# SexPeptID: an automated and reproducible workflow for paleoproteomics sex estimation in archaeological enamel

**DOI:** 10.64898/2026.06.05.730301

**Authors:** Marine Morvan

**Author notes:** Corresponding author (MM).

## Abstract

Accurate biological sex estimation is a key objective in archaeological and bioanthropological research but remains challenging when skeletal remains are fragmented, juvenile, or poorly preserved. Paleoproteomics approaches based on the detection of sex-specific amelogenin peptides (AMELX/AMELY) have emerged as a powerful alternative to osteological and genetic methods. However, current workflows often lack standardized criteria for peptide-level confidence assessment, potentially affecting the reproducibility and reliability of sex assignments.

In this study, I evaluated the impact of peptide-level confidence filtering on paleoproteomics-based sex estimation through the reanalysis of 164 *Homo sapiens* individuals from 10 published datasets and 26 *Bos taurus* individuals from 3 datasets, spanning contexts from the Pleistocene to the present. To address methodological inconsistencies, I developed SexPeptID, an R/Shiny-based framework that integrates Posterior Error Probability (PEP) filtering, standardized peptide selection, and explicit uncertainty assessment.

Application of SexPeptID revealed that peptide-level filtering substantially affects sex assignment outcomes: 17 previously classified males (10.4%) were reclassified as non-conclusive, while 5 individuals (3.1%) were identified as potentially female. Despite this sensitivity, AMELX/AMELY-based sex estimation remained robust overall, with stable signal ratios observed across archaeological periods. Variability in peptide intensities was primarily associated with dataset-specific factors rather than temporal differences, highlighting the influence of analytical workflows and preservation conditions.

By incorporating confidence-based filtering and a non-conclusive classification category, SexPeptID improves the transparency, reproducibility, and reliability of palaeoproteomics sex estimation, providing a standardized framework for future archaeological and bioanthropological studies.

**Highlights:**

- SexPeptID provides a reproducible framework for amelogenin-based sex estimation.
- Peptide-level confidence filtering significantly affects paleoproteomics sex estimates.
- 13.4% of published male assignments were revised after confidence filtering.
- AMELX/AMELY ratios show temporal stability from modern to Pleistocene samples.
- Standardized uncertainty assessment strengthens palaeoproteomics inference.

## 1. Introduction

Estimating biological sex is a fundamental aspect of anthropological, archaeological, and forensic research, providing essential information for reconstructing past populations and interpreting demographic and social structures (Mikšík et al., 2023). Traditionally, sex estimation relies on skeletal sexual dimorphism, particularly of the pelvis and skull (Bruzek and Murail, 2006). Osteomorphological methods are accurate in well-preserved adults but perform poorly in fragmented, degraded, or juvenile remains. Inter-population variability further limits the applicability of reference standards and increases classification uncertainty (Kotěrová et al., 2017). Sexual dimorphism in the skeleton arises from complex interactions among genetic background, hormonal regulation, environmental factors, and developmental processes (Dunsworth, 2020). Despite advances in statistical and bioinformatic methods, osteological sex estimation remains susceptible to misclassification (Waltenberger et al., 2025), motivating the development of molecular approaches for improved accuracy in challenging archaeological contexts.

Ancient genomics provides a complementary approach, particularly for incomplete or juvenile remains (Quincey et al., 2013; Stone et al., 1996). By analyzing molecular sexual dimorphism preserved in ancient DNA (aDNA), genomic methods can overcome morphological limitations and resolve osteological misclassifications (Ivarsson□Aalders et al., 2025). Approaches include shotgun-sequencing read-ratio methods, RY/RX ratios, and targeted PCR assays, often focusing on the amelogenin gene (AMELX/AMELY) (Praveen Kumar and Aswath, 2016). Detection of AMELX gene alone indicates a female, whereas the presence of both isoforms indicates a male. Additional strategies, such as Y-STR profiling and targeted capture enrichment, improve reliability in low-coverage or degraded datasets (Dash et al., 2020). Despite their utility, genomic methods face intrinsic limitations: aDNA is often highly degraded (<0.5% endogenous DNA in heavily degraded remains (Mittnik et al., 2016)), contamination is common, and alignment issues between X and Y regions, as well as structural complexity of the Y chromosome, can compromise accuracy. Rare Y-chromosome deletions may also produce erroneous female assignments (Guimarães et al., 2025).

Dental enamel is exceptionally well-preserved and provides an alternative substrate for sex estimation (Buonasera et al., 2020; Mikšík et al., 2023). Paleoproteomics approaches exploit molecular sexual dimorphism in amelogenin proteins, distinguishing AMELX and AMELY isoforms (Parker et al., 2019). Because amelogenin is restricted to enamel, this method reduces contamination from modern handlers. Proteins are also more resistant to degradation than aDNA, enabling reliable sex estimation even in poorly preserved samples. Paleoproteomics has been successfully applied to humans (adults and juveniles) (Brůžek et al., 2024; Drtikolová Kaupová et al., 2024; Lugli et al., 2019; Rebay-Salisbury et al., 2020), rare hominins (*Australopithecus africanus, Paranthropus robustus, Homo antecessor, Homo sp. Denisova*) (Demeter et al., 2022; Madupe et al., 2025b, 2025a; Tsutaya et al., 2025; Welker et al., 2020), and diverse mammals, including bovines, caprines, suids, mammoths, and rhinocerotids (Berezina et al., 2024; Blacka et al., 2025; Bray et al., 2025; Buckley et al., 2024; Cappellini et al., 2019; Green et al., 2019; Kotli et al., 2025, 2024; Rey-Iglesia et al., 2025). High-throughput LC-HRMS and MALDI-MS workflows allow sex estimation across large archaeological assemblages (Bray et al., 2026; Koenig et al., 2024). However, data interpretation remains manual, time-consuming, and dependent on bioinformatic expertise. Existing software was not designed for archaeological sex estimation, limiting reproducibility, scalability, and standardization.

Despite recent advances in paleoproteomics, sex assignment in ancient remains is still largely based on the detection of sex-specific peptides from amelogenin proteins (AMELX and AMELY), with limited standardization in peptide-level quality control. In many studies, peptide identifications are considered without explicit filtering based on statistical confidence metrics such as posterior error probability (PEP), potentially introducing variability in downstream interpretations. However, the absence of standardized peptide-level validation and confidence filtering limits reproducibility and comparability across studies. I hypothesize that peptide-level confidence filtering significantly affects sex assignment outcomes, particularly in low-quality archaeological proteomes.

To address this gap, I developed SexPeptID, an automated workflow for untargeted paleoproteomics sex estimation. SexPeptID provides an automated and reproducible framework to standardize peptide-level filtering, quantify uncertainty, and assess the robustness of proteomics sex estimation across heterogeneous datasets. The framework applies a default peptide-level confidence filter (PEP ≤ 0.05) but allows users to manually adjust thresholds, balancing reproducibility with analytical flexibility. This pipeline enables systematic assessment of peptide-level confidence on sex assignment, distinguishing robust biological signals from dataset- or quality-driven variability.

## 2. Materials and Methods

### 2.1. Materials

Paleoproteomics analyses were conducted using MaxQuant (version ≥2.7.0.0), a widely used software for mass spectrometry data processing. MaxQuant supports raw files from multiple instruments, including Thermo Fisher Scientific™ (.raw), mzML (.mzML), mzXML (.mzxml), and SCIEX WIFF (.wiff). Raw files from Agilent (.d), Bruker (.d), LECO (.peg), Shimadzu (.lcd), and Waters (*.raw) instruments can be converted to mzML or mzXML formats using tools such as MSConvert (Adusumilli and Mallick, 2017) or OpenMS (Röst et al., 2016) prior to analysis.

For this study, a custom FASTA database was built using the *Homo sapiens* AMELX (Q99217) and AMELY (Q99218) or *Bos taurus* AMELX (P02817) and AMELY (Q99004) sequences obtained from UniProt. Blank and replicate raw files were excluded. MaxQuant was run with the following recommended parameters: minimum peptide length of 6 amino acids, unspecific digestion (default), or the enzyme specified in the original sample preparation protocol (e.g., trypsin), variable modifications including deamidation (NQ) and oxidation (MP), a protein-level false discovery rate (FDR) of 1%, a minimum peptide score of 40 for both modified and unmodified peptides, and activation of dependent peptide identification and de novo sequencing.

### 2.2. Datasets Used

The following publicly available paleoproteomics datasets (10 for *Homo sapiens* and 3 for *Bos taurus*) were reanalyzed in this study.

### 2.3. Software and study design

All analyses were performed in R (version 4.5.1) using SexPeptID (https://github.com/MarineMorvan/SexPeptID), a custom Shiny application developed for reproducible peptide-level sex assignment. Data manipulation was carried out using dplyr, visualization with ggplot2, interactive tables with DT, and fragment ion annotation with ggrepel.

This study constitutes a reanalysis of publicly available paleoproteomics datasets. Its primary goal was to perform a sensitivity analysis assessing the impact of peptide-level confidence filtering on sex assignments. Unlike the original studies, which generally included all identified peptides regardless of Posterior Error Probability (PEP), SexPeptID provides explicit filtering and interactive visualization to evaluate the robustness of sex estimation.

SexPeptID can directly process MaxQuant output files (peptides.txt and msms.txt) without prior pre-processing, facilitating reproducible peptide selection, filtering, and downstream visualization.

### 2.4. Protein targets and species models

Sex estimation relied on amelogenin peptides encoded by X- and Y-chromosome homologs. Two species-specific models were implemented:

- Homo sapiens & hominins: AMELX (Q99217), AMELY (Q99218)
- Bos taurus: AMELX (P02817), AMELY (Q99004)

Selection of the species model determined the diagnostic peptide positions and rules for excluding ambiguous peptides. Peptides mapping to both X and Y isoforms were excluded to prevent misassignment.

### 2.5. Peptide selection and filtering

Peptides were extracted from MaxQuant peptides.txt using protein accession, peptide sequence, and start/end positions. Sex-dependent regions from different amelogenin isoforms were defined as:

- Homo sapiens & hominins: AMELX 28–29 (Q99217-2), 44–45 (Q99217-1), 58– 59(Q99217-3); AMELY 45 (Q99218-1), 59 (Q99218-1)
- Bos taurus: AMELX and AMELY 44, 48 (P02817, Q99004)

Only peptides overlapping these diagnostic sites were retained.

A peptide-level confidence filter based on Posterior Error Probability (PEP) was applied. A default PEP threshold (≤ 0.05) was applied, consistent with established paleoproteomics confidence standards, while allowing user adjustment for sensitivity analysis.

### 2.6. MS/MS integration and peptide-level quantification

Filtered peptides were merged with MaxQuant msms.txt to retrieve precursor intensities, PEP values, and fragment ion annotations. Fragment ion coverage was quantified by counting matched *b*- and *y*-ions and normalized by peptide length to account for sequence variability.

Peptide precursor intensities were summed per protein (AMELX and AMELY) per raw file. When multiple peptide-spectrum matches corresponded to the same peptide sequence, intensities were aggregated. Only peptides flagged as “included” in the SexPeptID interface (default: PEP ≤ 0.05) were used for downstream quantification, ensuring reproducibility and transparency in peptide selection.

### 2.7. Intensity processing and sex estimation

Zero intensities were replaced with ε = 1 prior to log transformation:

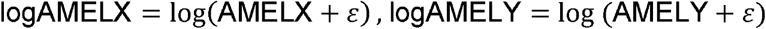

Male probability was estimated using a Y-chromosome intensity-based model:

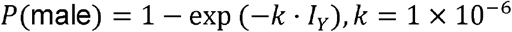

The parameter k was empirically defined based on observed AMELY intensity distributions to ensure conservative probability estimates and avoid overconfidence at low signal levels. This exponential formulation reflects diminishing returns in probability gain with increasing AMELY intensity.

### 2.9. Sex classification criteria

Individuals were classified into three categories:

- Female: no AMELY peptides detected
- Male: P(male) > 0.9
- Non-conclusive: all remaining cases, including:
  ∘ presence of AMELY peptides not included after PEP filtering
  ∘ insufficient peptide-level confidence
  ∘ low or inconsistent intensity signals

The non-conclusive category specifically captures uncertainty introduced by peptide-level filtering or sample degradation.

### 2.9. Data visualization

SexPeptID generated scatterplots of logAMELX versus logAMELY, with points colored by inferred sex and sample labels interactively displayed. Linear regression with 95% confidence intervals was applied to male samples where appropriate.

### 2.10. Output and reproducibility

SexPeptID outputs included:

- Summary tables with Raw file ID, logAMELX, logAMELY, P(male), and sex classification
- Publication-quality figures (TIFF, 12 × 8 inches)
- Interactive tables for peptide-level inspection and manual curation

The framework ensures reproducible peptide selection, transparent filtering decisions, and standardized sex classification across heterogeneous paleoproteomics datasets.

All analyses are fully reproducible using the same MaxQuant outputs and parameter settings within SexPeptID.

## 3. Results

### 3.1. Homo sapiens

In this study, I applied SexPeptID, a reproducible, statistically grounded filtering and decision framework based on peptide-level PEP and intensity-derived signals, to reassess published sex assignments of 164 individuals from 10 public datasets spanning the modern period to the Pleistocene. This analysis can be viewed as a sensitivity assessment of published sex assignments to peptide-level confidence filtering, as original studies typically included all identified peptides regardless of confidence metrics such as PEP. Across all datasets, SexPeptID reclassified 17 males (10.4%) as non-conclusives and 5 males (3.1%) as potentially females (Figure 1). Notably, dataset PXD026419 accounted for 5 of the non-conclusives reclassifications of males and all cases of males reclassified as potentially females (Supplementary Material S1). These results demonstrate that proteomics sex assignment is sensitive to peptide-level confidence filtering, particularly in datasets exhibiting higher signal variability.

**Figure 1:**
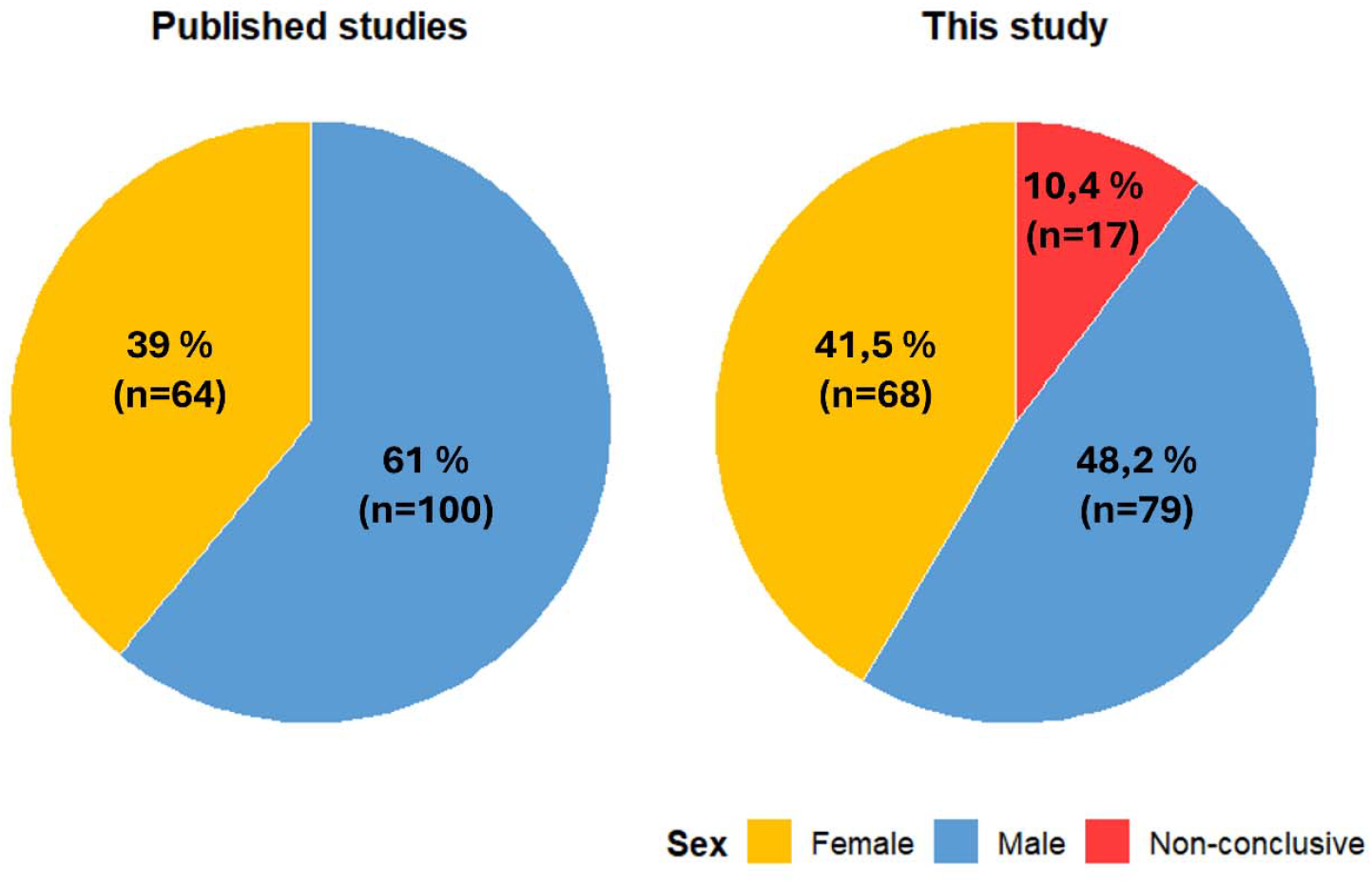
Impact of peptide-level confidence filtering on sex assignment across datasets.

The distribution of log_10_(AMELX/AMELY) ratios among individuals classified as male was examined across archaeological periods (Figure 2), with some periods combining multiple datasets. Overall, the ratio remained remarkably stable over time. Although slight differences in median values were observed between periods, they remained within the same order of magnitude, and no significant temporal variation was detected (Kruskal–Wallis test, *p* = 0.075). The Neolithic period, represented by a single individual (*n* = 1), was not considered interpretable.

**Figure 2:**
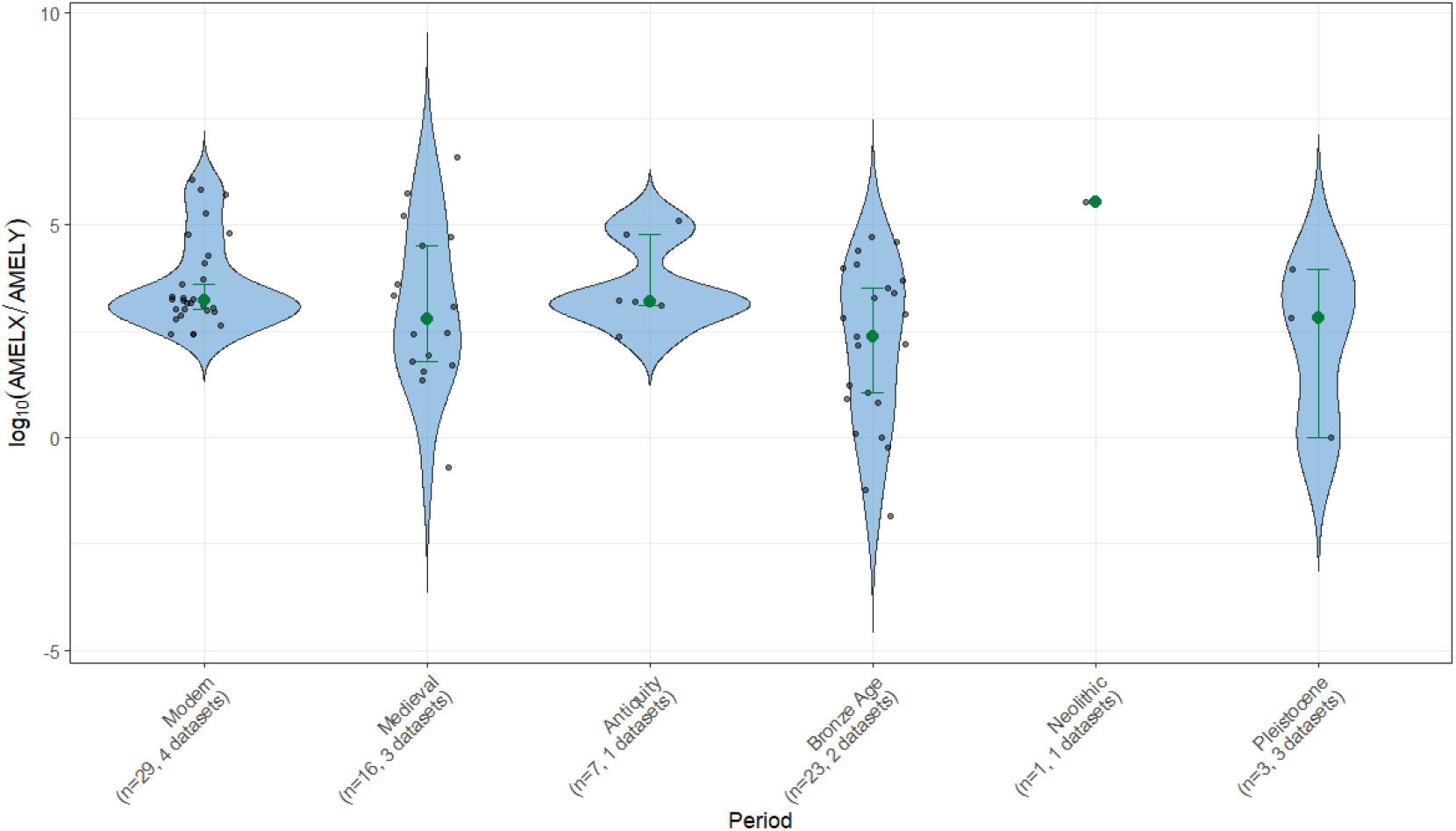
Distribution of log_10_(AMELX/AMELY) ratios among male individuals across archaeological periods.

These findings indicate that the AMELX/AMELY signal ratio is generally stable across archaeological periods in male individuals. The consistent co-detection of AMELX and AMELY peptides suggests that paleoproteomics sex estimation is not substantially affected by temporal biases, and that the AMELX/AMELY report alone does not provide information to predict the archaeological period of an unknown sample. However, ratios falling within the range observed in male individuals may provide additional support for male sex assignment. Notably, this temporal stability was observed after applying peptide-level filtering with SexPeptID to datasets originating from multiple studies and generated across different laboratories and mass spectrometry platforms (Table 1), further supporting the robustness of the underlying biological signal.

**Table 1:**
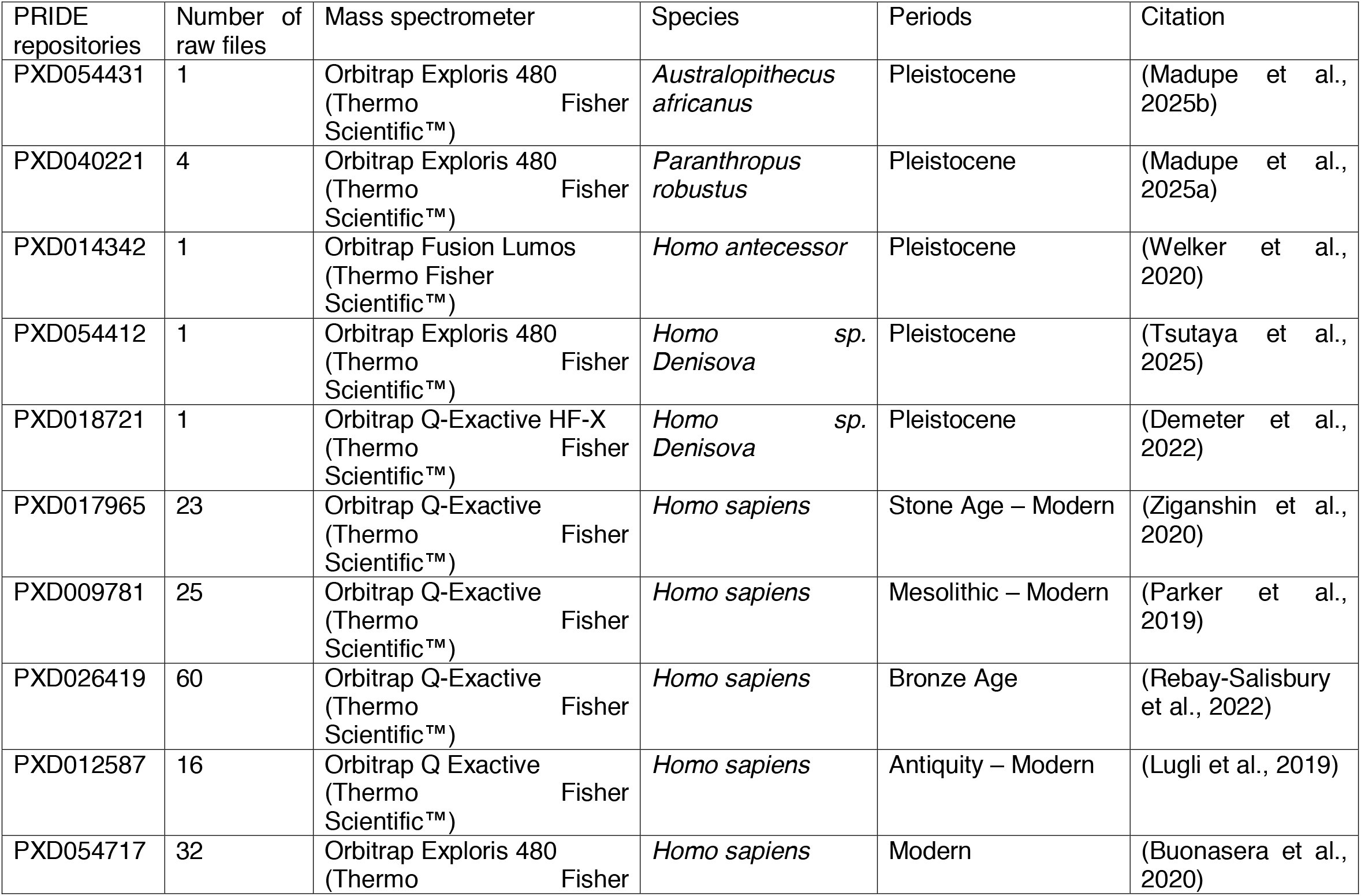
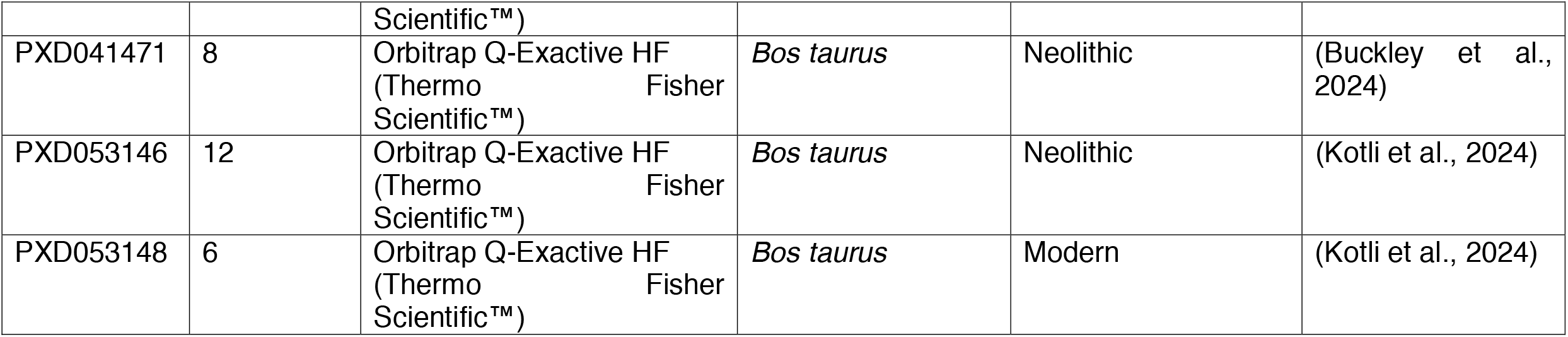
Publicly available paleoproteomics datasets reanalyzed.

To further investigate inter-study variability, log_10_(AMELX/AMELY) ratios were plotted separately for each dataset (Figure S1). While several datasets show tight clustering around their median, others exhibit broader dispersion. In particular, dataset PXD026419 displays a markedly wider distribution compared to most other datasets. Despite this variability, median values within datasets generally remain consistent with the overall trend observed across archaeological periods. These findings suggest that variability in log-ratios is primarily structured at the dataset level rather than the temporal level. Accordingly, differences between studies, including analytical workflows and sample-specific factors, appear to contribute more to the observed dispersion than archaeological period differences. This further highlight that part of the observed heterogeneity across studies may reflect methodological sensitivity to peptide inclusion criteria.

I next examined AMELX and AMELY peptide intensities across individuals, stratified according to the sex assignments generated by SexPeptID (Figure 3). The resulting boxplots provide an overview of the distribution of these markers in relation to sex and sample quality. AMELX intensities are broadly similar between males and females, a pattern consistent with dosage-compensation mechanisms associated with X-chromosome inactivation (Heard and Disteche, 2006), which limit expression differences between XX and XY individuals. In contrast, individuals classified as non-conclusive exhibit substantially lower AMELX signals. AMELX does not discriminate sex but serves as a proxy for overall proteome preservation. AMELY, by contrast, shows a clearly sex-specific pattern. Male individuals consistently display strong AMELY signals, whereas non-conclusive samples often show weak or undetectable AMELY. AMELY is a sex-specific marker reliably detected in males when peptide recovery is sufficient. Together, these results emphasize the complementary roles of AMELX and AMELY. AMELX primarily reflects sample quality, while AMELY provides the sex-specific signal required for confident male identification, explaining the robustness of SexPeptID across a wide range of archaeological contexts. Overall, this supports the idea that stricter peptide-level confidence filtering primarily affects low-quality spectra rather than the underlying biological signal.

**Figure 3:**
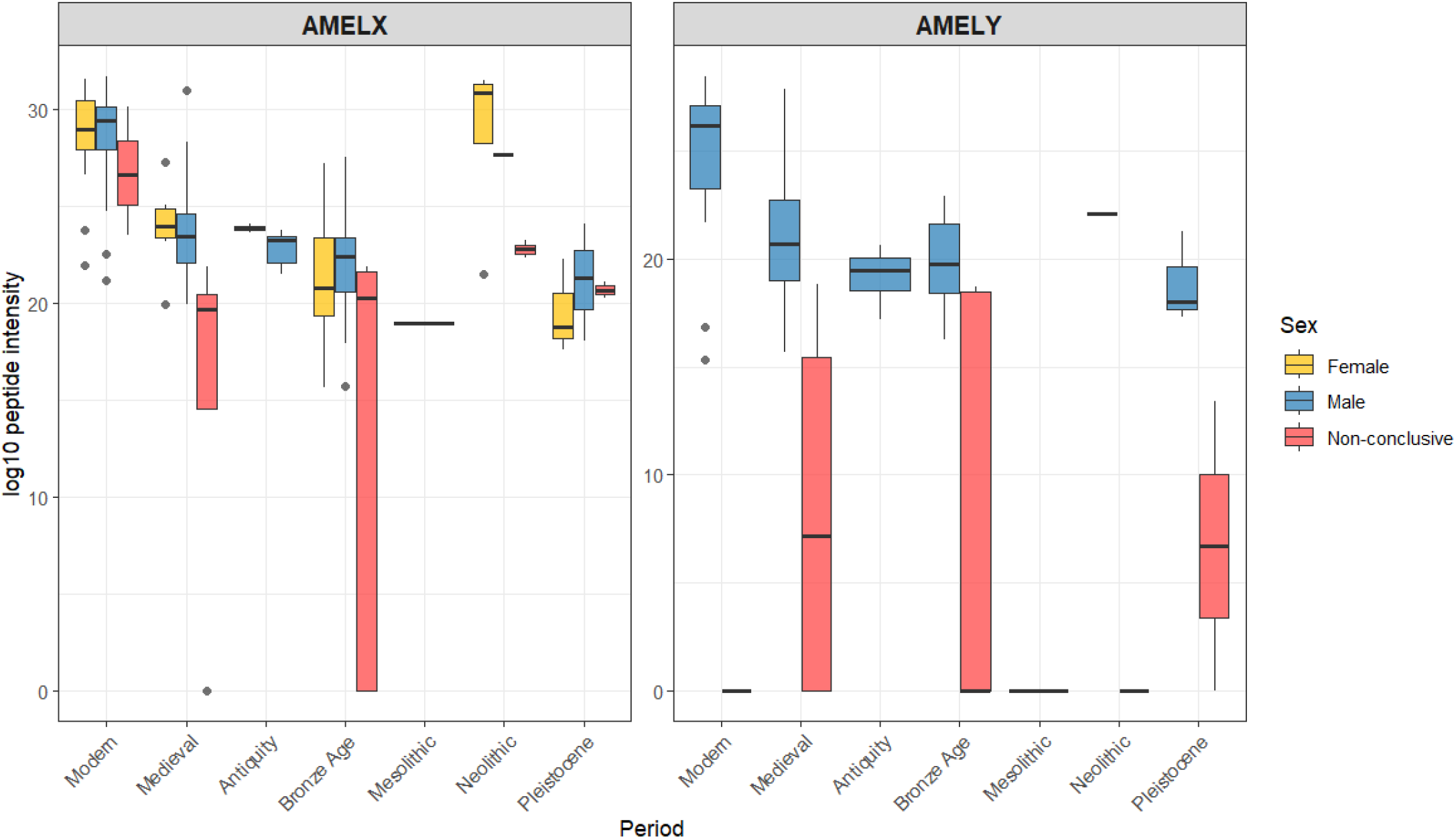
Distribution of AMELX and AMELY intensities according to sex classification.

### 3.2. Multiple species integration

The distribution of log_10_(AMELX/AMELY) ratios among *Homo sapiens* and *Bos taurus* individuals classified as male was examined across archaeological periods (Figure 4), with some periods combining multiple datasets. A significant difference is observed between Modern and Neolithic in *Bos taurus* males (Kruskal-Wallis, *p* = 0.0208), but the interpretation is limited by the small number of samples (n = 26) and the reduced temporal coverage (n = 2).

**Figure 4:**
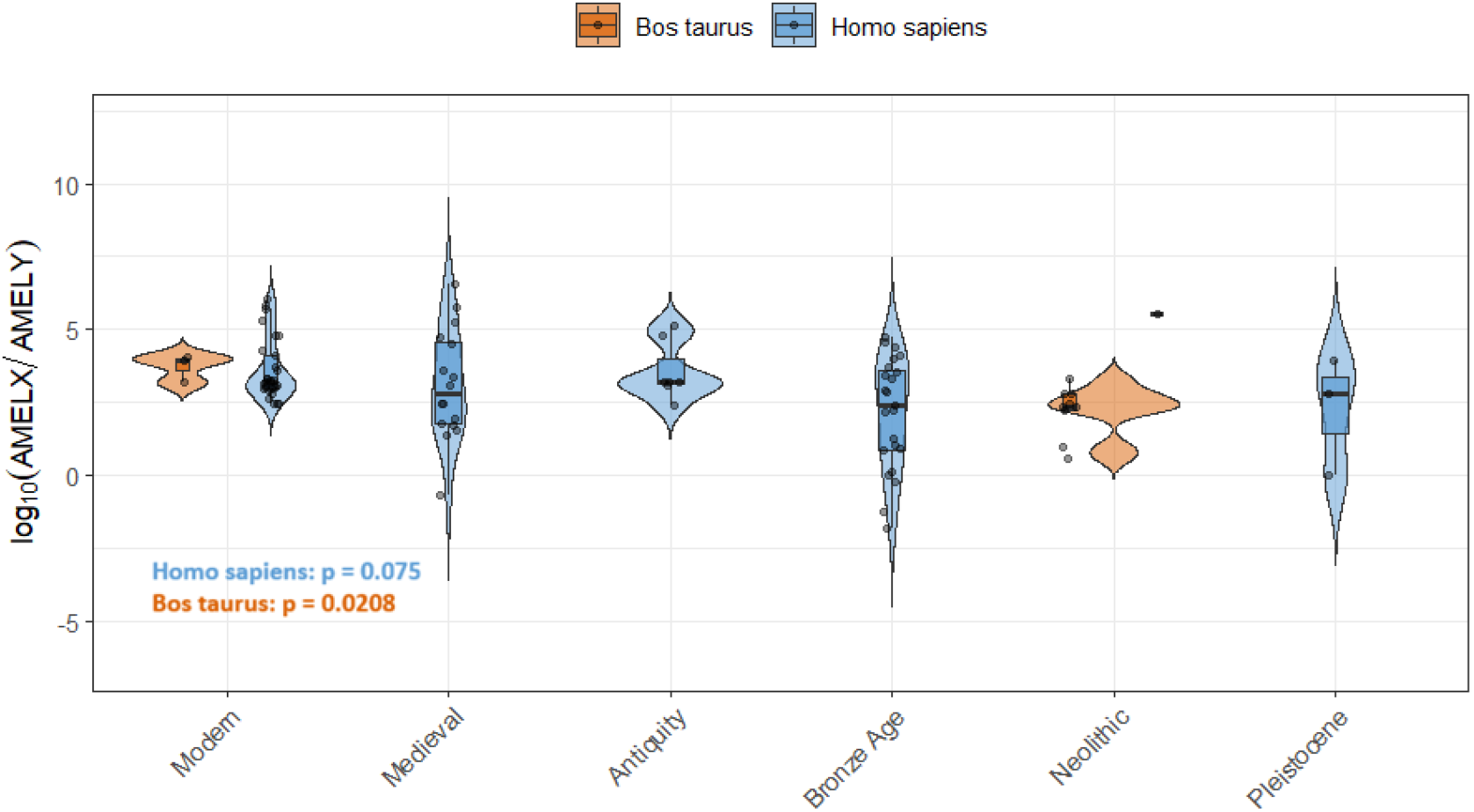
Distribution of *Homo sapiens* and *Bos taurus* log_10_(AMELX/AMELY) ratios among male individuals across archaeological periods.

## 4. Discussion

These results demonstrate that peptide-level confidence filtering is a major determinant of sex assignment outcomes in archaeological proteomics. By reanalyzing 164 *Homo sapiens* individuals and 26 *Bos taurus* individuals, I show that applying standardized filtering criteria leads to the reclassification of a subset of previously assigned individuals, highlighting the importance of peptide-level validation.

### 4.1. Robustness and reproducibility of SexPeptID

SexPeptID combines peptide-level PEP with summed precursor intensities, allowing the classification of samples into male, female, or non-conclusive categories. Most previous studies have relied on the presence or absence of AMELY peptides without explicit peptide-level confidence filtering. Unlike the original publications, which included all peptides regardless of confidence, my analyses demonstrate that a small but meaningful fraction of previously classified males (10.4%) become non-conclusive, and a subset (3.1%) may be reclassified as potentially female. These findings confirm that peptide-level confidence is a critical determinant of reliable proteomics sex assignment.

The interface in Shiny ensures reproducibility while remaining user-friendly. Users can directly input raw MaxQuant files without pre-processing, and peptides with PEP > 0.05 are automatically deselected, though manual inclusion is possible. This design facilitates both routine and exploratory analyses, making SexPeptID broadly applicable for palaeoproteomic datasets.

### 4.2. Temporal stability of AMELX/AMELY signals

Across archaeological periods, log_10_(AMELX/AMELY) ratios in males remained remarkably consistent for *Homo sapiens*, with no evident temporal trend. This suggests that the combined detection of AMELX and AMELY peptides is robust over time, supporting the use of this ratio as a reliable marker for sex rather than for chronological inference.

Dataset-level variability was observed, particularly in PXD026419, but median values remained consistent with overall trends. These findings indicate that inter-study differences (e.g., sample preparation, MS workflows, preservation conditions) contribute more to variation than the archaeological period itself.

Regarding *Bos taurus*, despite the limited number of modern male samples analyzed (n = 6), the median log_10_(AMELX/AMELY) ratio was higher than that observed in *Homo sapiens* males (n = 29). Further investigation using a larger dataset encompassing samples from different chronological periods is required to confirm this pattern. If validated, species-specific differences in AMELX/AMELY ratios could provide a useful criterion for distinguishing between species and for assigning unknown samples to their species of origin.

### 4.3. Complementary roles of AMELX and AMELY

This intensity-based analysis reinforces the complementary nature of these markers:

- AMELX intensity largely reflects overall proteome recovery and is similar across males and females. Lower AMELX intensities correlate with non-conclusive samples, indicating that AMELX is a robust quality indicator rather than a sex-specific marker.
- AMELY intensity is distinctly sex-specific, reliably detected in males when peptide recovery is sufficient. Samples with weak or undetectable AMELY signals are classified as non-conclusive, highlighting the importance of peptide-level confidence filtering for accurate male assignment.

Together, these observations explain the robust performance of SexPeptID across heterogeneous datasets and preservation contexts.

### 4.4. Implications for palaeoproteomics and archaeology

This work provides a systematic framework to:

1. Evaluate the reliability of published sex assignments
2. Standardize sexing workflows across datasets and periods
3. Provide reproducible, transparent criteria for peptide inclusion

By explicitly accounting for peptide-level confidence, SexPeptID reduces the risk of misclassification due to low-quality or ambiguous peptides, which is particularly important for ancient samples where degradation is frequent.

Our study also demonstrates that a non-conclusive category is essential to reflect uncertainty realistically, avoiding overinterpretation of degraded or low-signal samples, a principle that should be adopted in future palaeoproteomic studies.

### 4.5. Limitations and future directions

- Although the analysis of 164 *Homo sapiens* individuals from 10 datasets provides a solid foundation for evaluating the method, increasing sample sizes would improve the robustness of statistical inferences, especially for chronologically underrepresented periods. Furthermore, the incorporation of larger datasets from additional species will be essential to determine whether interspecific differences can be reliably detected and to assess the feasibility of species identification based on these patterns.
- These results extend previous observations on proteome degradation by explicitly quantifying its impact on sex assignment outcomes.
- This study relies on MaxQuant-derived peptide identifications, and results may vary with alternative search engines or parameter settings.

## 5. Conclusion

SexPeptID provides a reproducible and statistically grounded framework for palaeoproteomics sex estimation by integrating peptide-level confidence metrics into the sex assignment process. Through the reanalysis of 164 *Homo sapiens* individuals from 10 public datasets and 26 *Bos taurus* individuals, this study demonstrates that peptide-level filtering can substantially influence sex assignment outcomes, leading to the reclassification of 17 previously identified males as non-conclusive and 5 as potentially female. These results highlight the importance of explicitly accounting for peptide confidence when interpreting AMELX and AMELY evidence.

At the same time, the analyses confirm the overall robustness of AMELX/AMELY-based sex estimation across a broad temporal range, from modern to Pleistocene contexts. The stability of AMELX/AMELY ratios among male individuals suggests that the underlying biological signal remains largely preserved despite differences in archaeological age, while observed variability is primarily associated with dataset-specific and methodological factors.

By combining standardized peptide filtering, intensity-based evaluation, and an explicit non-conclusive category, SexPeptID improves the transparency, reproducibility, and reliability of paleoproteomics-based sex estimation. Beyond reassessing published assignments, the framework provides a practical tool for harmonizing analyses across studies and laboratories, facilitating more robust comparisons between datasets. As palaeoproteomics continues to expand, approaches that formally integrate uncertainty and quality assessment, such as SexPeptID, will be essential for strengthening bioarchaeological inference and ensuring the reproducibility of future research.

## CRediT authorship contribution statement

Conceptualization: MM, Data curation: MM, Formal analysis: MM, Funding acquisition: MM, Investigation: MM, Methodology: MM, Project administration: MM, Resources: MM, Software: MM, Validation: MM, Visualization: MM, Writing – original draft: MM, Writing – review & editing: MM

## Data Availability Statement

The custom R/Shiny-based workflow (SexPeptID) used for peptide-level filtering and sex assignment is publicly available on GitHub at: https://github.com/MarineMorvan/SexPeptID. All analyses were performed using publicly available paleoproteomics datasets retrieved from the PRIDE repository (accession numbers provided in Table 1).

## Acknowledgements

I warmly thanked all authors who shared their paleoproteomics datasets.

## Appendix A. Supplementary data

Supplementary data to this article can be found online at

