## Supplementary material for "SexPeptID: an automated and reproducible workflow for paleoproteomics sex estimation in archaeological enamel": Figure S1

Marine Morvan^1,*^

^1^ Vilnius University, Faculty of History, Department of Archaeology, Universiteto g. 7, LT-01513 Vilnius, Lithuania

ORCID MM: 0000-0001-7904-7108

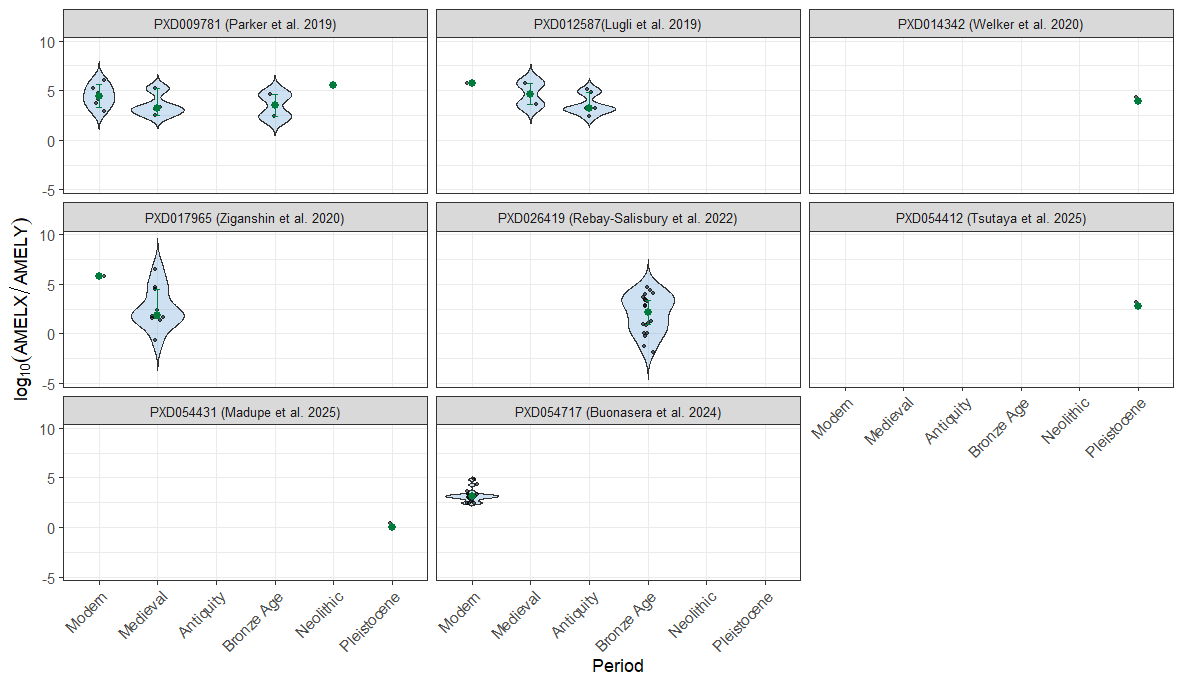

Figure S1: Distribution of log_10_(AMELX/AMELY) ratios among male individuals across archaeological periods according to each dataset.
